# Cell wall damage impairs root hair cell patterning and tissue morphogenesis mediated by the Arabidopsis receptor kinase STRUBBELIG

**DOI:** 10.1101/2020.11.04.368241

**Authors:** Ajeet Chaudhary, Xia Chen, Barbara Leśniewska, Jin Gao, Sebastian Wolf, Kay Schneitz

## Abstract

Cell wall remodeling is essential for the control of growth and development as well as the regulation of stress responses. However, the underlying cell wall monitoring mechanisms remain poorly understood. Regulation of root hair fate and flower development in *Arabidopsis thaliana* requires signaling mediated by the atypical receptor kinase STRUBBELIG (SUB). Furthermore, SUB is involved in cell wall integrity signaling and regulates the cellular response to reduced levels of cellulose, a central component of the cell wall. Here, we show that continuous exposure to sub-lethal doses of the cellulose biosynthesis inhibitor isoxaben results in altered root hair patterning and floral morphogenesis. Genetically impairing cellulose biosynthesis also results in root hair patterning defects. We further show that isoxaben exerts its developmental effects through the attenuation of SUB signaling. Our evidence indicates that down-regulation of SUB is a multi-step process and involves changes in SUB complex architecture at the plasma membrane, enhanced removal of SUB from the cell surface, and downregulation of *SUB* transcript levels. The results provide molecular insight into how the cell wall regulates cell fate and tissue morphogenesis.

## Introduction

Plant cells are encapsulated by a semi-rigid cell wall. As a consequence, cell wall remodeling represents a central pillar in the control of growth, development, and the defense against abiotic and biotic stresses. In recent years, the mechanisms monitoring and modulating cell wall integrity in plants have received much attention (1–3). However, the molecular framework controlling signaling from the cell wall during development and stress responses is still poorly understood.

The plant cell wall is composed of carbohydrates, including cellulose, hemicellulose, pectin, and phenolic compounds, but also contains a large number of cell-wall-bound proteins (4, 5). Cellulose synthesis is carried out by cellulose synthase (CESA) protein complexes (CSCs) at the plasma membrane. The CESA1, CESA3 and CESA6 subunits represent the core CSC subunits of the primary cell wall of expanding cells (6, 7). The herbicide isoxaben induces a rapid clearing of CESA complexes from the plasma membrane (8). The reaction to cellulose biosynthesis inhibition (CBI) induced by isoxaben or by defects in CESA subunits represents a well-characterized compensatory cell wall damage (CWD) response (3, 9). Factors implied as cell wall sensors mediating this response include the cell surface receptor kinase THESEUS 1 (THE1) (10), a member of the Catharanthus roseus RECEPTOR-LIKE KINASE 1-LIKE (CrRLK1L) subfamily, and the leucine-rich repeat (LRR) receptor kinase MIK2/LRR-KISS (11), among others (2).

The role of the cell wall in morphogenesis has long been appreciated. Cell wall remodeling is central to cellular growth (12). In addition, the cell wall tightly connects neighboring cells. There is growing evidence that differential growth in physically coupled cells cause mechanical stresses that may in turn influence morphogenesis (13–15). Interestingly, the extracellular matrix of animal cells as well as the cellulose-containing cell wall of brown algae has been shown to influence cell fate (16–18) but a clear demonstration of a similar role for the plant cell wall is missing.

Cell fate regulation in Arabidopsis involves the atypical leucine-rich repeat receptor kinase STRUBBELIG (SUB). SUB, also known as SCRAMBLED (SCM), controls root hair specification (19, 20). SUB regulates additional developmental processes, including floral morphogenesis and integument outgrowth (21, 22). Present evidence indicates that SUB fulfills its role in these developmental processes in a complex with the transmembrane protein QUIRKY (QKY) (20, 23–25). Recent work revealed that SUB also participates in the isoxaben-induced CWD response in young seedlings (26). Interestingly, QKY was found to play only a minor role in this process. Moreover, SUB, THE1, and MIK2 appear to function in different CBI-induced CWD pathways (26).

Here, we report that SUB activity is regulated by the cell wall. We show that exposing plants to sub-lethal doses of isoxaben results in sub-like morphological defects. We also show that cell wall alterations eventually cause altered SUB complex architecture, increased endocytosis of SUB, and reduced SUB transcript levels. Our data further reveal that ectopic upregulation of SUB expression counteracts the assayed morphological effects of cellulose deficiency.

## Results

### Isoxaben reduces *SUB* expression levels

In light of the role of *SUB* in the CBI-induced cell wall damage response (26) we investigated if isoxaben modulates *SUB* activity. To this end we made use of a well-characterized line carrying the *sub-1* null allele and a fully complementing transgene encoding a SUB:EGFP translational fusion driven by its endogenous promoter (pSUB::SUB:EGFP) (24, 27, 28). We noticed considerably weaker pSUB::SUB:EGFP reporter signal in liquid-grown seedlings exposed to 600 nM isoxaben in comparison to reporter signal in mock-treated seedlings (Fig. 1A,B). Signal reduction could be detected from around five hours after the start of the treatment and was clearly evident after eight hours. Signal did not completely disappear but reached about 50 percent of the intensity detected prior to the start of the isoxaben treatment. Importantly, reporter signal strength appeared unaltered in isoxaben-resistant *ixr2-1* seedlings carrying a mutation in the *CESA6* gene (Fig. S1) (29–31).

**Fig. 1.**
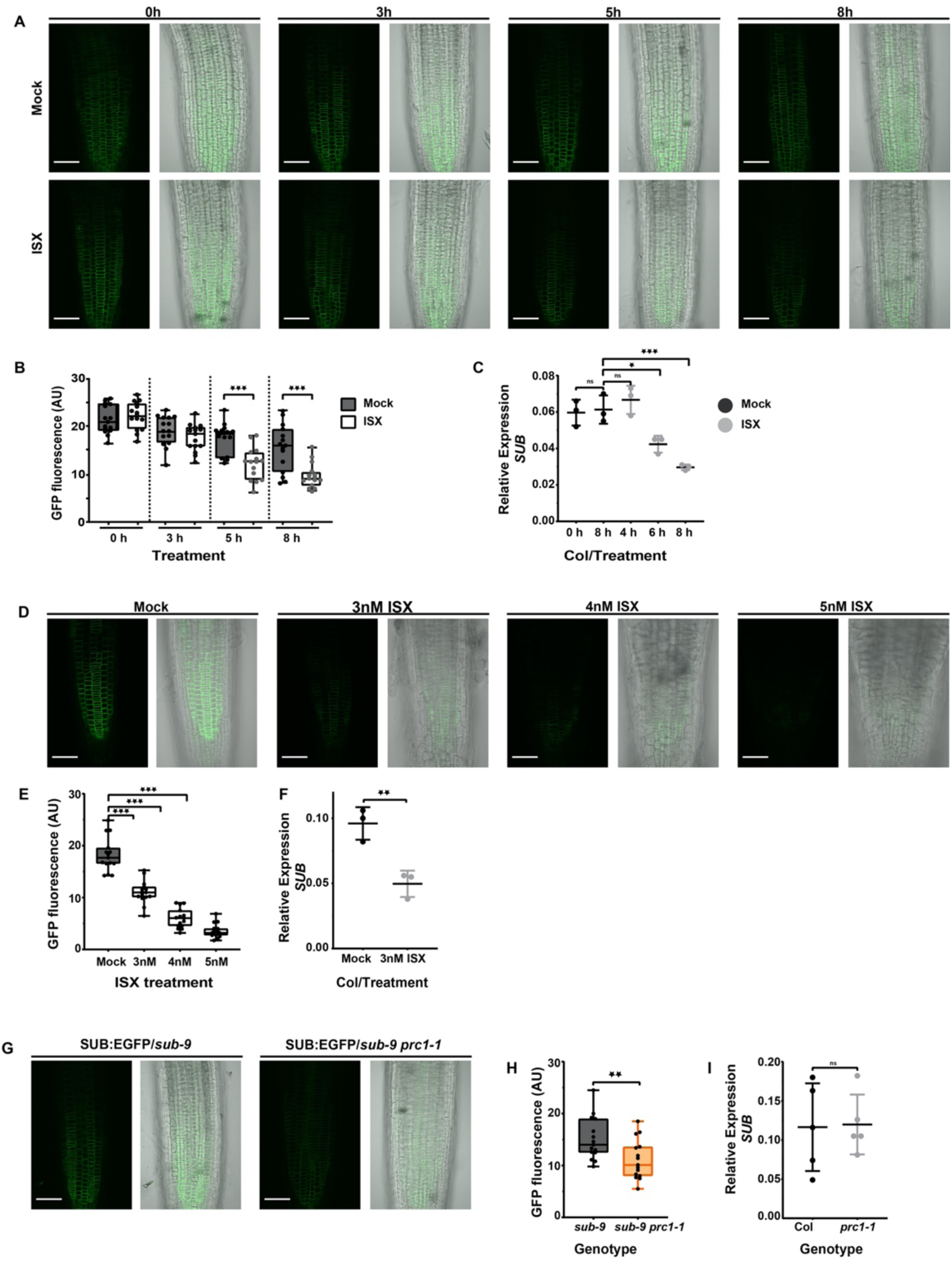
Effect of isoxaben treatment on *SUB* expression. (A), (D) and (G) show confocal micrographs depicting optical sections through the roots of seven-day-old seedlings. (A) Signal intensity of a functional pSUB::SUB:EGFP reporter upon exposure of liquid-grown seedlings to mock or 600 nM isoxaben. Duration of treatment and treatment are indicated. Imaging parameters between the mock and isoxaben treatments were identical. Note the reduction of signal in isoxaben-treated seedlings that starts at about five hours of treatment. (B) Quantification of the data shown in (A). Box and whiskers plots are shown. n =15. Asterisks represent statistical significance (***, P < 0.0006; one-way ANOVA followed by post hoc Tukey’s multiple comparison test). The experiment was performed three times with similar results. (C) Relative mRNA levels of *SUB* in seven days-old seedlings exposed to mock or 600 nM isoxaben for eight hours. Expression was detected by qPCR. Mean ± SD is shown, n = 3 biological replicates each with mean of three technical replicates. Asterisks represent statistical significance (*, P < 0.02; ****, P < 0.0001; one-way ANOVA followed by post hoc Tukey’s multiple comparison test). The experiment was performed three times with similar results. (D) Signal intensity of a functional pSUB::SUB:EGFP reporter upon exposure of plate-grown seedlings continuously exposed to mock or three, four, or five nM isoxaben. Duration of treatment and treatment are indicated. Imaging parameters between the mock and isoxaben treatments were identical. Note the reduced signal in isoxaben-treated seedlings. (E) Quantification of the data shown in (D). Box and whiskers plots are shown. N =15. Asterisks represent statistical significance (****, P < 0.0001; one-way ANOVA followed by post hoc Tukey’s multiple comparison test). The experiment was performed three times with similar results. (F) Relative expression of *SUB* in seven days-old seedlings grown on plates containing mock or 3 nM isoxaben. Expression was detected by qPCR. Mean ± SD is shown, n = 3 biological replicates each with mean of three technical replicates. Asterisks represent statistical significance (**, P < 0.009; unpaired t-test with Welch’s correction, two-tailed P values). (G) Signal intensity of a functional pSUB::SUB:EGFP reporter in seven day old *sub-9* and *sub-9 prc1-1* plate-grown seedlings. Imaging parameters between both genotype were identical. Note the reduced signal in *sub-9 prc1-1* seedlings. (H) Quantification of the data shown in (D). Box and whiskers blots are shown. n ≤ 15. Asterisks represent statistical significance (**, P=0.0101; unpaired t-test with Welch’s correction, two-tailed P values). The experiment was performed three times with similar results. (I) Relative expression of *SUB* in seven days-old seedlings grown on plates in Col and *prc1-1*. Expression was detected by qPCR. Mean ± SD is shown, n = 3 biological replicates each with mean of three technical replicates. Asterisks represent statistical significance (ns= not significant; unpaired t-test with Welch’s correction, two-tailed P values). The experiment was performed three times with similar results. Scale bars: 50 μm.

Furthermore, we detected significantly diminished endogenous SUB mRNA levels in seedlings treated with isoxaben for up to eight hours by quantitative real-time polymerase chain reaction (qPCR) in five out of six biological replicates (Fig. 1C). Transcript levels were noticeably reduced already at the six hour time point in three out of three biological replicates.

Next, we tested the effects of prolonged exposure of seedlings to isoxaben on *SUB* activity. Arabidopsis seedlings growing in presence of isoxaben exhibit a response gamut ranging from near normal growth to essentially no growth in the narrow range of 1 to 10 nM isoxaben, with an I50 at 4.5 nM (32). Thus, we chose to analyze seven-days-old *sub-1 pSUB::SUB:EGFP* seedlings that were continuously grown on agar plates containing 3 nM, 4 nM, and 5 nM isoxaben, respectively. In comparison to mock treated samples we observed a concentration-dependent decrease in reporter signal in isoxaben-treated seedlings (Fig. 1D,E). We then assessed if endogenous *SUB* transcript levels were affected in plate-grown wild-type seedlings exposed to 3 nM isoxaben for seven days. We detected a reduction of *SUB* transcript levels by about 50 percent in comparison to untreated seedlings (Fig. 1F).

*PRC1* encodes the CESA6 subunit of cellulose synthase and the predicted null allele *prc1-1* shows reduced cellulose levels (33). To assess if SUB activity is also diminished when cellulose biosynthesis is genetically perturbed we generated a *sub-9 prc1-1* double mutant homozygous for the pSUB::SUB:EGFP reporter. We then compared the reporter signal in root tips of seven days-old plate-grown seedlings of the strong *sub-9* mutant and the *sub-9 prc1-1* double mutant. We observed a noticeable reduction in reporter signal in *sub-9 prc1-1* in comparison to *sub-9* (Fig. 1G). Next, we analyzed endogenous *SUB* transcript levels in wild-type and *prc1-1* seedlings by qPCR but we did not detect significant differences between the two genotypes (Fig. 1H).

We then investigated if downregulation of *SUB* by isoxaben involves *QKY* function. To this end we generated *pSUB::SUB:EGFP qky-8* plants and analyzed the signal of the pSUB::SUB:EGFP reporter in the roots of the respective plate-grown seedlings. We observed an additive effect on reporter signal strength when comparing untreated seedlings with seedlings exposed to isoxaben (Fig. S2). The result indicates that isoxaben and *QKY* affect SUB abundance through parallel pathways in seedling roots.

### Cellulose biosynthesis inhibition affects SUB-complex architecture at the plasma membrane

The results mentioned above prompted us to investigate if isoxaben influences the composition of SUB-containing protein complexes at the plasma membrane. To this end we assessed the steady-state fluorescence anisotropy of SUB:EGFP following mock or isoxaben treatment. Fluorescence anisotropy (*r*) describes the rotational freedom of a fluorescent molecule, such as GFP. Upon protein homo-oligomerization of GFP-based fusion proteins Förster resonance energy transfer (FRET) can occur (homo-FRET) resulting in a decrease in fluorescence anisotropy (34–36). Monitoring changes in fluorescence anisotropy has been successfully applied in receptor kinase interaction studies involving for example CLV1 or BAK1 (37, 38). As control we used two lines carrying translational fusions of the TARGET OF MP5 (TMO7) transcription factor to one or three GFP moieties, respectively (39). We measured the fluorescence anisotropy values for the two fusion proteins in the nuclei of epidermal cells of the root meristem of plate-grown seedlings. We observed a fluorescence anisotropy of 0.38 for TMO7:1xGFP and 0.26 for TMO7:3xGFP (Fig. 2A,B,F). Free GFP in a plant cell has a steady-state anisotropy value of 0.33 (37, 38). The higher anisotropy of TMO7:1xGFP indicates that this fusion protein is more restricted in its rotational freedom than free GFP. The low value for TMO7:3xGFP is indicative of homo-FRET between the three closely-linked GFP moieties.

**Fig. 2.**
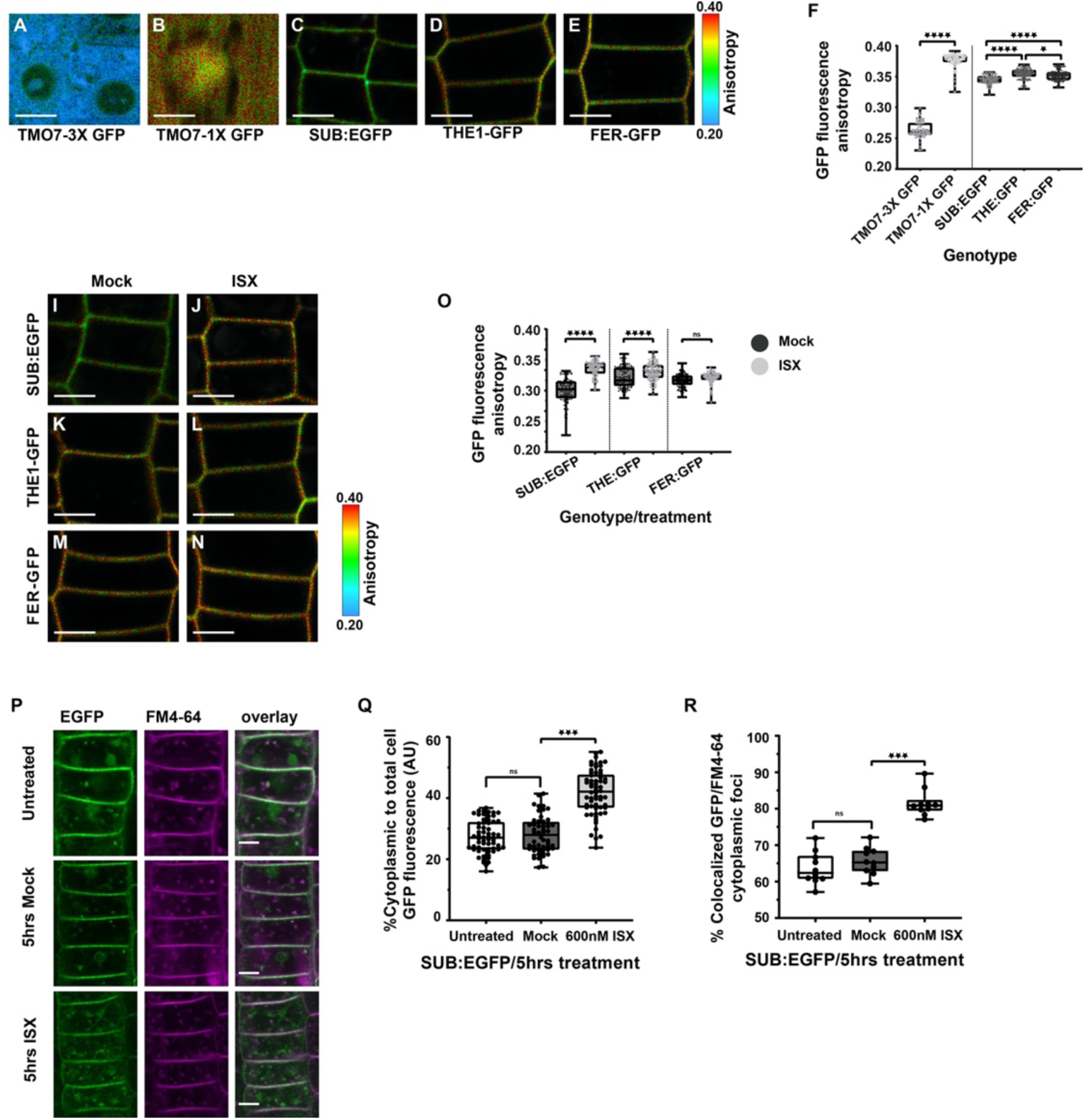
Effect of isoxaben treatment on fluorescence anisotropy and sub-cellular localization of pSUB::SUB:EGFP. (A-E) show confocal micrographs depicting fluorescence anisotropy of GFP in root epidermal cells of the meristematic region in seven-day-old seedlings. (A,B) Fluorescence anisotropy of TMO7:3xGFP and TMO7:1xGFP in cytoplasm. Note the increased anisotropy in TMO7:1xGFP (B). (C, D) and (E) depict fluorescence anisotropy at the plasma membrane of the functional reporters pSUB::SUB:EGFP, pTHE1::THE1:GFP and pFER::FER:GFP, respectively. Imaging and fluorescence anisotropy calculation parameters were identical. A minimum of 25 photons per anisotropy event were counted. (F) Quantification of the data shown in (A) to (E). Box and whisker plots are shown. 150 ≤ n ≤ 169. Asterisks represent statistical significance (*, P < 0.04; ****, P < 0.0001; one-way ANOVA followed by Sidak’s multiple comparison test). The experiment was performed three times with similar results. (G-L) show confocal micrographs depicting GFP-based fluorescence anisotropy in root epidermal cells of the meristematic region in seven-day-old seedlings after 4.5 hours mock or 600nM isoxaben treatment. Genotype and treatment is indicated. Imaging and fluorescence anisotropy calculation parameters were identical. A minimum of 25 photons per anisotropy event were counted. Note the increased anisotropy after isoxaben treatment in SUB:EGFP (G,H) and THE1::GFP (I,J) respectively. (M) Quantification of the data shown in (G) to (L). Box and whisker plots are shown. 121 ≤ n ≤ 185. Asterisks represent statistical significance (****, P < 0.0001; one-way ANOVA followed by Sidak’s multiple comparison test). The experiment was performed three times with similar results. (N) Fluorescence micrographs show optical sections of epidermal cells of root meristem of seven-day-old seedlings. Duration of treatment and treatment are indicated. Endocytic vesicles are marked by FM4-64. Note the increased SUB:EGFP signal in the cytoplasm as well as increased SUB:EGFP signal abundance in FM4-64 marked vesicles after isoxaben treatment. (O) Box and whisker plot depicting quantitative analysis of ratio of SUB:EGFP signal intensity in cytoplasm to total cell shown in (N). N is 52 ≤ n ≤ 61 cells from 10 different roots. Asterisks represent statistical significance (***, P < 0.0001; one way ANOVA followed by Sidak’s multiple comparison test). The experiment was performed three times with similar results. (P) Box and whisker plot depicts the result of co-localization analysis of SUB:EGFP and FM4-64 shown in (N). N=10 roots, each data point represents a mean of at least 5 cells per root. Asterisks represent statistical significance (***, P < 0.0001; one way ANOVA followed by Sidak’s multiple comparison test). The experiment was performed three times with similar results. Scale bars: 5 μm.

We then determined the fluorescence anisotropy values for SUB:EGFP localized at the plasma membrane of epidermal cells of the root meristem. In addition, we investigated two additional receptor kinases translationally fused to GFP and driven by their native promoters: THESEUS1 (THE1) and its homolog FERONIA (FER). THE1 and FER are both implied in monitoring the cell wall status (10, 40, 41). In cells of untreated plate-grown seedlings we observed a fluorescence anisotropy value of 0.345 for SUB:EGFP, 0.355 for THE1:GFP and 0.352 for FER:GFP (Fig. 2C-F). The results indicate that SUB:EGFP has a slightly higher rotational freedom in comparison to THE1:GFP and FER:GFP. We then analyzed the fluorescence anisotropy values of these reporters upon isoxaben treatment. To this end plate-grown seedlings were transferred onto MS plates containing 600 nM isoxaben and incubated for 4.5 hours. We observed a fluorescence anisotropy value for SUB:EGFP of 0.340 in mock-treated seedlings and a value of 0.361 upon isoxaben treatment. We scored fluorescence anisotropy values of 0.352 and 0.358 for THE1:GFP and 0.350 and 0.353 for FER:GFP, respectively. Thus, there is a noticeable difference in fluorescence anisotropy values for SUB:EGFP in mock versus isoxaben-treated cells. By contrast, we detected only minor alterations in these values for THE1:GFP and FER:GFP. The results indicate that SUB:EGFP-containing protein complexes experience different architectures depending on the presence or absence of isoxaben-induced CWD.

### Perturbation of cellulose biosynthesis promotes internalization of SUB

We then tested if isoxaben treatment affects the subcellular distribution of the pSUB::SUB:EGFP reporter signal in epidermal cells of the root meristem (Fig. 2N-P). In untreated seedlings SUB is known to undergo ubiquitination and continuous internalization (25, 28). We found that the percentage of cytoplasmic SUB:EGFP foci increased upon isoxaben treatment in comparison to untreated or mock-treated cells (Fig. 2N,O). To assess if the accumulation of cytoplasmic SUB:EGFP foci upon isoxaben treatment related to endocytosis we imaged cells of seedlings that were exposed to the different types of treatment upon a 5-min incubation with the endocytic tracer dye FM4-64. Applying a convenient criterion for colocalization (28, 42), the internal SUB:EGFP and FM4-64 signals were considered colocalized when the distance between the centers of the two types of signals was below the limit of resolution of the objective (0.24 μm). In untreated or mock-treated seedlings we observed that 63.7% (n = 322) and 65.6% (n = 536), respectively, of all cytoplasmic SUB:EGFP foci were also marked by FM4-64 confirming the previous observation that SUB:EGFP undergoes recognizable internalization in the absence of any apparent stimulation (28). Upon isoxaben treatment, we noticed that 81.5% (n=633) of all cytoplasmic SUB:EGFP foci were also marked by FM4-64. These data support the notion that isoxaben treatment eventually leads to increased endocytosis of SUB:EGFP.

### Exposing plants to sub-lethal doses of isoxaben induces root hair patterning defects

Since isoxaben treatment results in the downregulation of *SUB* we investigated if isoxaben treatment of wild-type plants results in a *sub*-like phenotype. In a first step we explored if application of isoxaben induces root hair patterning defects in seven-days-old seedlings. To this end, we compared the number of hair (H) and nonhair (N) cells in the N and H positions of the root epidermis, respectively, in untreated and treated wild type seedlings to the respective numbers in untreated *sub-9* roots (Fig. 3) (Table 1). In untreated plate-grown wild-type plants we found that 97.4 percent of cells at the H position were hair cells while only 1.9 percent of cells at the N position were hair cells (Fig.3A) (Table 1). In contrast, roots of wild-type seedlings grown on 3 nM isoxaben plates for seven days exhibited 67.7 percent hair cells in the H position and 24.4 percent hair cells in the N position (Fig. 3B). A similar value was observed for *sub-9* mutants, confirming previous results (19). Root hair patterning appeared unaltered in mock or isoxaben-treated *ixr2-1* mutants (Fig. 3D,E). To test if a genetic defect in cellulose biosynthesis results in aberrant root hair patterning we analyzed *prc1-1* mutants. We observed a mild but robust difference in root hair patterning in comparison to wild type (Fig. 3F,G).

**Table 1.** Distribution of root hair and nonhair cells in the root epidermis.

| Genotype | roots<sup>a</sup> | H position<sup>b</sup> |  | N position<sup>b</sup> |  |
| --- | --- | --- | --- | --- | --- |
|  |  | Hair (%) | Nonhair (%) | Hair (%) | Nonhair (%) |
| Col-0 | 17 | 97.4 ± 3.3 | 2.6 ± 3.3 | 1.9 ± 2.8 | 98.1 ± 2.8 |
| Col-0 / 3 nM ISX | 17 | 67.7 ± 4.9 | 32.3 ± 4.8 | 24.4 ± 6.0 | 75.6 ± 6.0 |
| *sub-9* | 17 | 70.9 ± 4.1 | 29.8 ± 4.1 | 23.4 ± 4.1 | 76.6 ± 4.1 |

<sup>a</sup>Total number of different roots analyzed.
<sup>b</sup>Mean ± standard deviation. Total number of cells counted at H position: 321 ≤ n ≤ 442. Total number of cells counted at N position: 472 ≤ n ≤ 561.
The experiment was repeated twice with similar results.

**Fig. 3.**
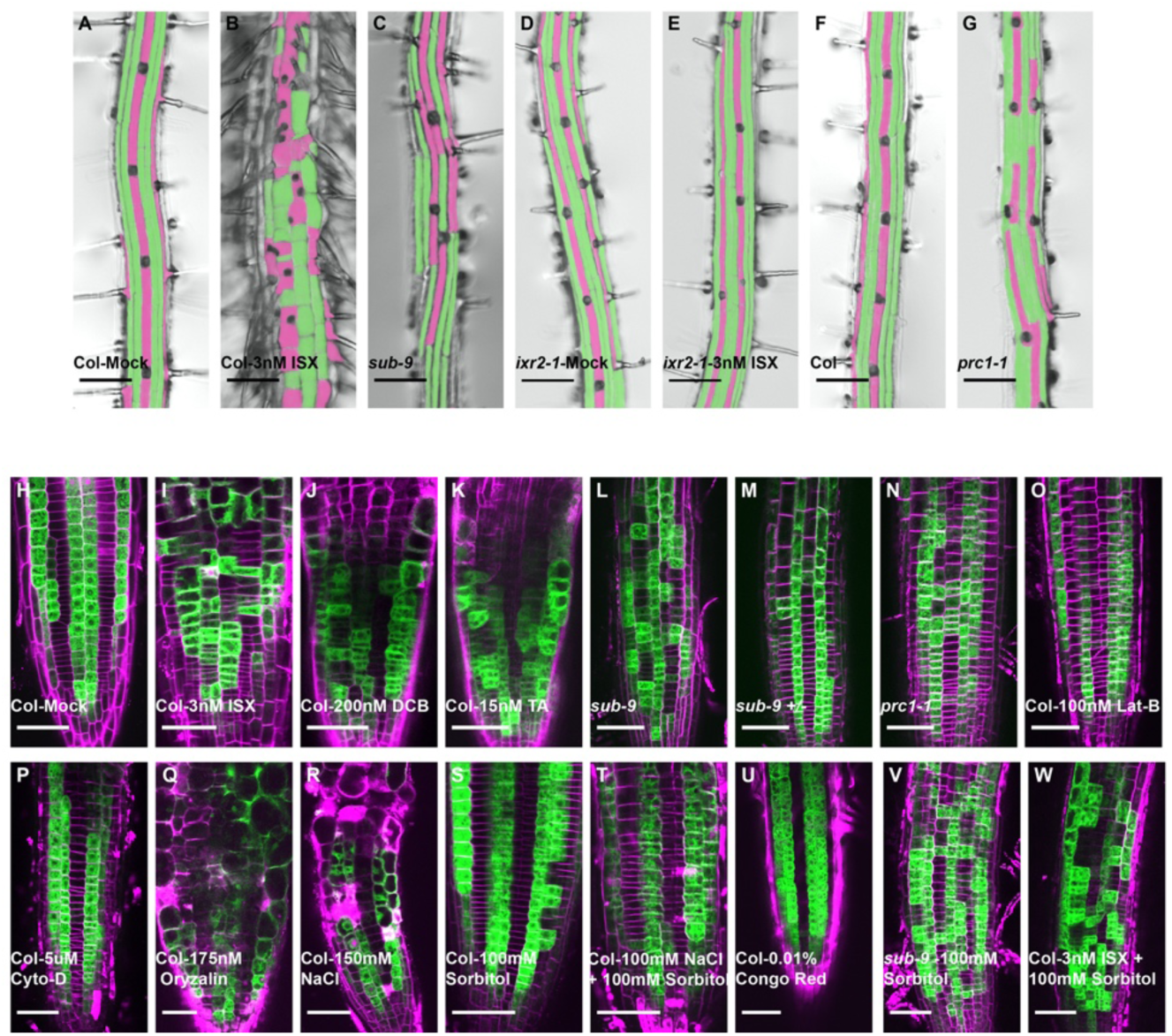
Reduced cellulose biosynthesis results in altered root hair cell patterning. (A-G) Micrographs show epidermal root hair cell patterning in seven-day-old seedlings in presence or absence of isoxaben. Genotypes and/or treatments are indicated. Pink and green colors mark root hair cells and non-root hair cells, respectively. Note the straight-line arrangement of hair cells. For a quantification, see Table 1. The experiment was performed three times with similar results. (H-W) Confocal micrographs depicting optical sections through roots of seven-day-old seedlings showing pGL2::GUS:EGFP reporter signal. The cell wall was counter-stained with propidium iodide, except for (U). Genotypes and/or treatments are indicated. For a quantification of some of the effects see Table 2. (H) All analyzed seedlings showed this pattern (20/20. (I) 26/29. (J) 25/28. (K) 27/27. (L) 25/25. (M) 26/38. (N) 23/33. (O) 27/33. (P) 32/38 (Q) 25/33 (R) 27/34 (S) 28/32 (T) 28/35 (U) 22/24 (V) 32/32 (W) 34/34. The remaining roots showed the occasional single cell displacement in the reporter expression as also sometimes seen in wild type (Table 2). The experiment was performed three times with similar results. Scale bars: (A-G) 100 μm, (H-W) 50 μm.

Next, we tested whether the altered root hair patterning is reflected at the molecular level. We made use of Col-0 plants carrying a pGL2::GUS:EGFP reporter. The *GLABRA2* (*GL2*) promoter drives expression in N cells but not in H cells of the root epidermis (43), and the expression pattern of a reporter driven by the *GL2* promoter thus serves as a convenient and faithful proxy for root hair patterning (19, 28, 44). Upon exposing wild-type seedlings first grown on MS plates for four days to 1 to 3 nM isoxaben for 48 hours we found a pronounced and concentration-dependent increase in defects in the expression pattern of the pGL2::GUS:EGFP reporter (Fig. Fig. 3H,I,) (Table 2). The prominent mis-expression resembled the expression pattern of the reporter in *sub-9* mutants (Fig. 3H,I,L,M) (Table 2).

**Table 2.** Position-dependent pattern of pGL2::GUS:EGFP reporter expression in root epidermal cells upon a 48 hour exposure to isoxaben.

| Genotype | ISX [nM]<sup>a</sup> | H cells total<sup>b</sup> | H Position<sup>c</sup> | N cells total<sup>b</sup> | N Position<sup>c</sup> |
| --- | --- | --- | --- | --- | --- |
| Col | 0 | 633 | 2.7 | 741 | 96.6 |
| *sub-9* | 0 | 548 | 33.6<sup>d</sup> | 615 | 74.8 |
| Col | 1 | 441 | 14.1<sup>e</sup> | 589 | 86.1 |
|  | 2 | 375 | 26.7 | 459 | 78.7 |
|  | 3 | 395 | 30.6 | 579 | 77.0 |
| *SUB OE L1* | 0 | 423 | 6.9 | 558 | 77.3 |
|  | 1 | 616 | 4.7<sup>f</sup> | 663 | 95.9 |
|  | 2 | 327 | 1.2<sup>g</sup> | 396 | 92.2 |
|  | 3 | 401 | 16.7 | 445 | 86.7 |
| *SUB OE O3* | 0 | 508 | 4.7 | 701 | 82.2 |
|  | 1 | 638 | 3.3<sup>h</sup> | 706 | 94.5 |
|  | 2 | 320 | 3.4<sup>i</sup> | 396 | 92.4 |
|  | 3 | 359 | 10.9 | 475 | 86.1 |

For each genotype and treatment at least 10 different roots of seven-days-old seedlings were analyzed.
<sup>a</sup>Isoxaben (ISX) concentration in plate
<sup>b</sup>Total number of cells at H or N position scored
<sup>c</sup>Percentage of cells at H or N position expressing the pGL2::GUS:EGFP reporter
<sup>d</sup>Statistical significance: *sub-9* vs Col (untreated), $P < 0.0001$ .
<sup>e</sup>Statistical significance: Col (1 nM ISX) vs Col (untreated), $P < 0.0001$ .
<sup>f</sup>Statistical significance: L1 (1 nM ISX) vs Col (untreated), $P = 0.0708$ ; L1 (1 nM ISX) vs Col (1 nM ISX), $P < 0.0001$ . L1 (1 nM ISX) vs L1 (untreated), $P = 0.1686$ .
<sup>g</sup>Statistical significance: L1 (2 nM ISX) vs Col (untreated), $P = 0.1676$ ; L1 (2 nM ISX) vs Col (2 nM ISX), $P < 0.0001$ , L1 (2 nM ISX) vs L1 (untreated), $P = 0.0001$ .
<sup>h</sup>Statistical significance: O3 (1 nM ISX) vs Col (untreated), $P = 0.6219$ ; O3 (1 nM ISX) vs Col (1 nM ISX), $P < 0.0001$ ; O3 (1 nM ISX) vs O3 (untreated), $P = 0.2241$ .

We similarly treated wild-type seedlings with 200 nM 2,6-dichlorobenzonitrile (DCB) or 15 nM Thaxtomin A, two other cellulose biosynthesis inhibitors (45–47). Both drugs induced *sub*-like expression pattern defects of the pGL2::GUS:EGFP reporter (Fig. 3J,K). Moreover, we observed that *prc1-1* mutants exhibited a mild but noticeable aberration in the expression pattern of the reporter (Fig. 3N). Thus, plants treated with three different cellulose inhibitors and a cellulose biosynthesis mutant all exhibit similar defects in expression of the pGL2::GUS:EGFP reporter.

We then explored if drugs affecting the cytoskeleton had an effect on the pGL2::GUS:EGFP expression pattern. We grew seedlings for four days on MS plates followed by exposure for 48 hours to 100 nM latrunculin B, 5μM Cytochalasin D, or 175 nM oryzalin, respectively. However, in all instances we failed to notice an obvious effect on the pGL2::GUS:EGFP expression pattern (Fig. 3O-Q) although the applied concentrations affect the organization of microtubules and actin filaments (48–50). We also tested if application of a cell wall stain has an effect on the pGL2::GUS:EGFP pattern. We did not observe a noticeable effect upon staining with Congo Red, a stain that allows the detection of cellulose microfibrils (51) (Fig. 3U).

The isoxaben-induced CWD response is generally sensitive to turgor pressure (52). To investigate if the effect of isoxaben on root hair patterning follows the same pattern we tested pGL2::GUS:EGFP expression in sorbitol-treated *sub-9* seedings and in wild-type seedlings simultaneously exposed to isoxaben and sorbitol. In both instances we found the pGL2::GUS:EGFP expression pattern to be aberrant (Fig. 3V,W) indicating that the isoxaben-induced effect on root hair patterning is not sensitive to alterations in turgor pressure. Thus, these data indicate that the CWD-sensitive mechanism controlling *SUB* activity is distinct from the regulation of other CBI-induced CWD responses, including callose accumulation, cell cycle gene expression, or root cell shape changes (52–54).

### Exposing plants to sub-lethal doses of isoxaben induces *sub*-like floral defects

To explore further the effect of isoxaben on tissue morphogenesis we tested if isoxaben treatment also induces *sub*-like defects in flowers and ovules. To this end, we compared wild-type (L*er*) and *sub-1* plants that were cultivated on soil in the presence of isoxaben. Plants were initially grown without any treatment. Just before bolting we began watering wild-type plants with 100 to 500 nM isoxaben and continued watering with isoxaben in three-day intervals for two weeks. We then analyzed stage 3 floral meristems (stages according to (55) (Fig. 4A-C). Floral meristems of *sub-1* mutants show aberrant cell division planes in the L2 layer (20, 21). Analysis of floral meristems of isoxaben-treated wild-type Col-0 plants revealed similar defects (Fig. 4B) the frequency of which increased with increasing concentrations of isoxaben (Table 3). Next, we compared stage 13 flowers of isoxaben-treated L*er* plants to flowers from untreated *sub-1* plants (Fig. 4D-F). We noticed that flowers from isoxaben-treated plants exhibited a *sub*-like altered arrangement of petals. Finally, we analyzed the ovule phenotype of isoxaben-treated wild-type plants. We noticed *sub*-like defects in integument outgrowth in late stage 3 or early stage 4 ovules (ovule stages according to (56)) (Fig. 4G-I) (Table 4). The frequency and severeness of the integument defects also depended on the concentration of isoxaben (Table 4).

**Table 3.** Number of periclinal cell divisions in L2 layer of stage 3 floral meristems of plants exposed to different concentrations of isoxaben.

| Genotype | ISX [nM]<sup>a</sup> | N FM<sup>a</sup> | N PCD<sup>b</sup> | Percentage |
| --- | --- | --- | --- | --- |
| Col | 0 | 29 | 5 | 17.2 |
| *sub-9* | 0 | 31 | 14 | 45.2 |
| Col | 200 | 14 | 4 | 28.6 |
|  | 300 | 21 | 8 | 38.1 |
|  | 400 | 25 | 15 | 60.0 |
| *SUB OE L1* | 0 | 8 | 2 | 25.0 |
|  | 400 | 27 | 6 | 22.2 |

<sup>a</sup>Number of floral meristems analyzed.
<sup>b</sup>Number of periclinal cell divisions observed.
The experiment was performed twice with similar results.

**Table 4.**
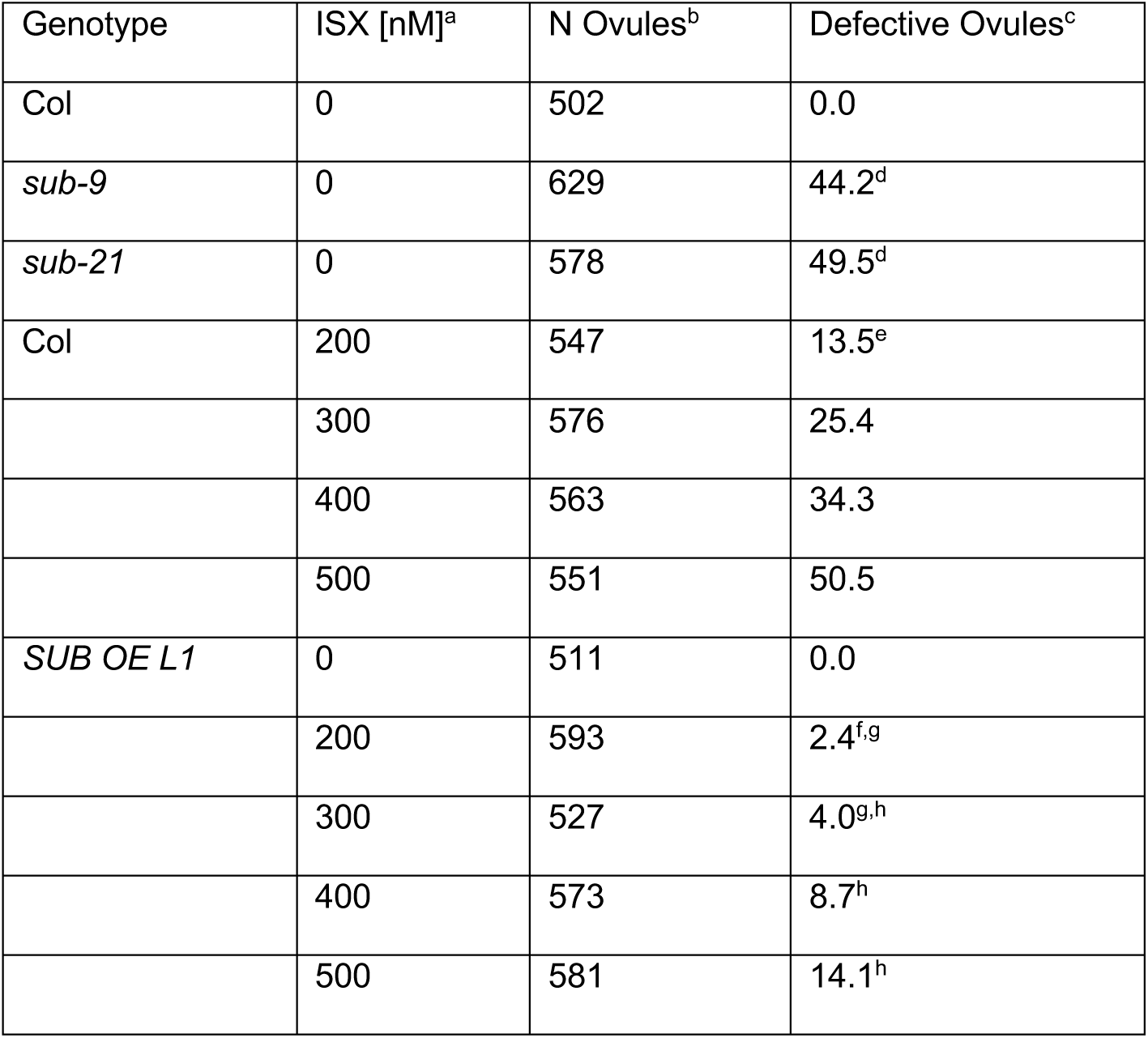
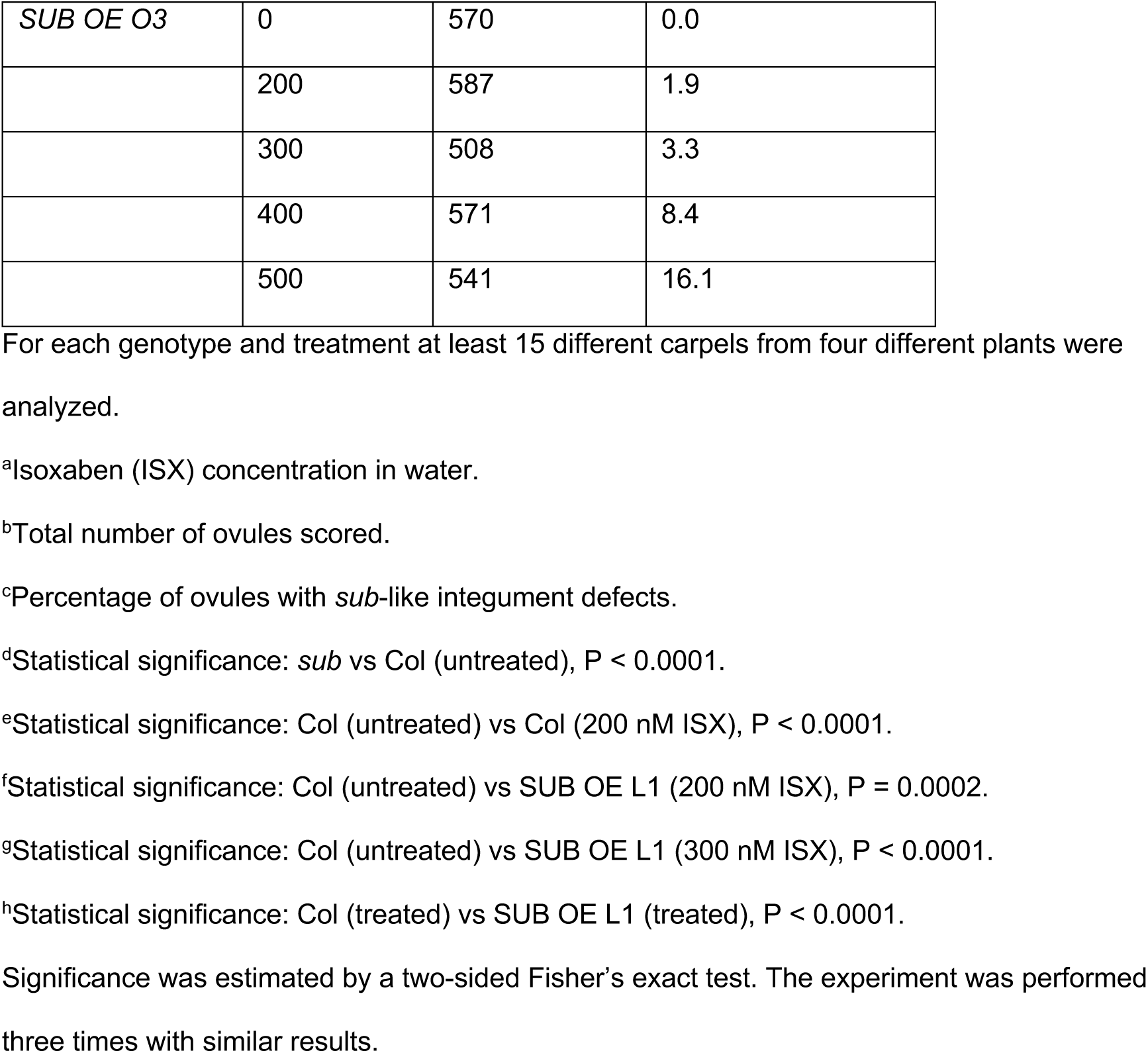
Comparison of integument defects between Col, *sub-9, sub-21*, and Col plants exposed to different concentrations of isoxaben.

**Fig. 4.**
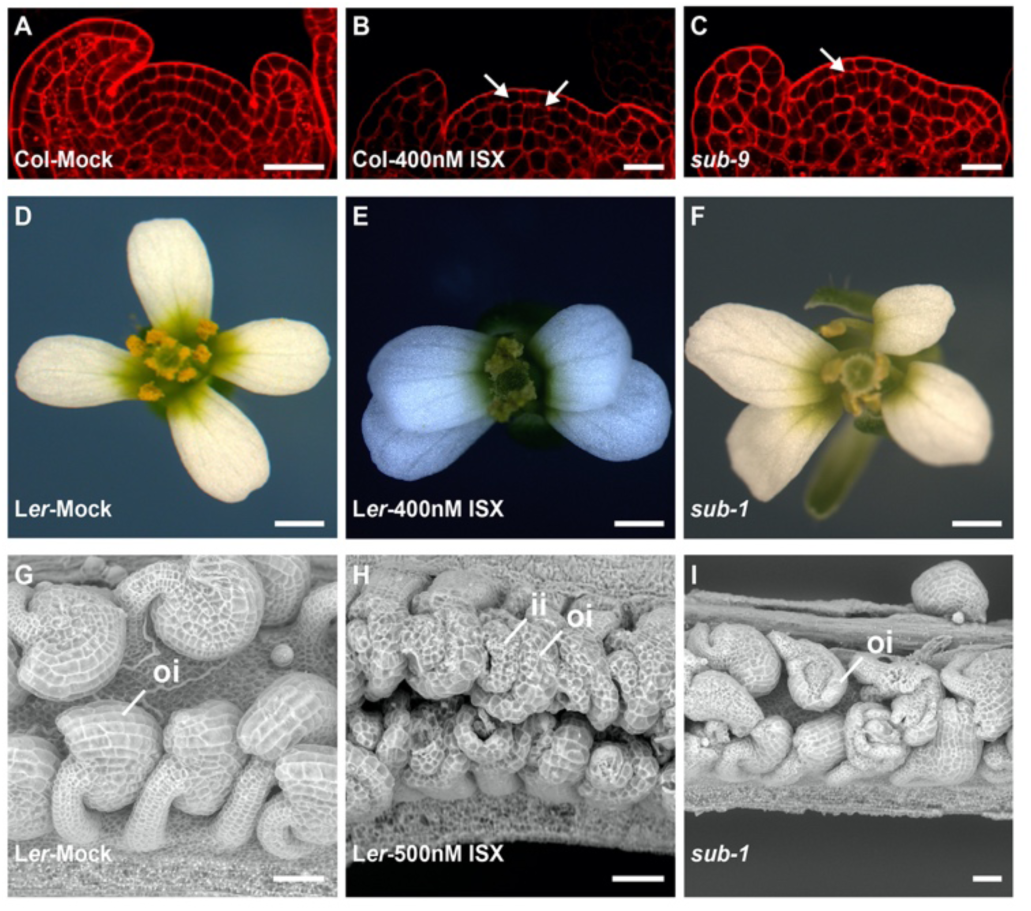
Effect of isoxaben treatment on floral and ovule morphogenesis. Genotypes and treatments are indicated. For a quantification of the effects see Table 3. (A-C) Confocal micrographs depicting mid-optical sections through stage 3 floral meristems. Arrows indicate irregular periclinal cell divisions in L2. (D-F) Morphology of mature stage 13 or 14 flowers. (E,F) Compare to (D). Note the aberrant arrangement of petals. (G-I) Electron scanning micrographs depicting mature stage 3 or 4 ovules. (H,I) Compare to (G). Note aberrant integuments. Scale bars: (A-C) 20 μm; (D-F) 0.5 mm; (G-I) 50 μm.

### Ectopic expression of *SUB* attenuates the detrimental effects of isoxaben on root hair patterning and floral development

If isoxaben treatment results in a downregulation of *SUB* and a *sub*-like phenocopy ectopic expression of *SUB* should counteract this outcome. We tested this hypothesis by analyzing the effects of isoxaben on two well-characterized lines carrying a pUBQ::SUB:mCherry transgene (lines L1 and O3) (26). To this end we generated L1 and O3 lines homozygous for the pGL2::GUS:EGFP construct. We then analyzed reporter signal in seven-days-old plate-grown seedlings that had been grown on normal plates for five days before being transferred to plates containing no isoxaben or 1 nM, 2 nM, or 3 nM isoxaben, respectively, for another 48 hours prior to analysis (Fig. 5A-D) (Table 2).

**Fig. 5.**
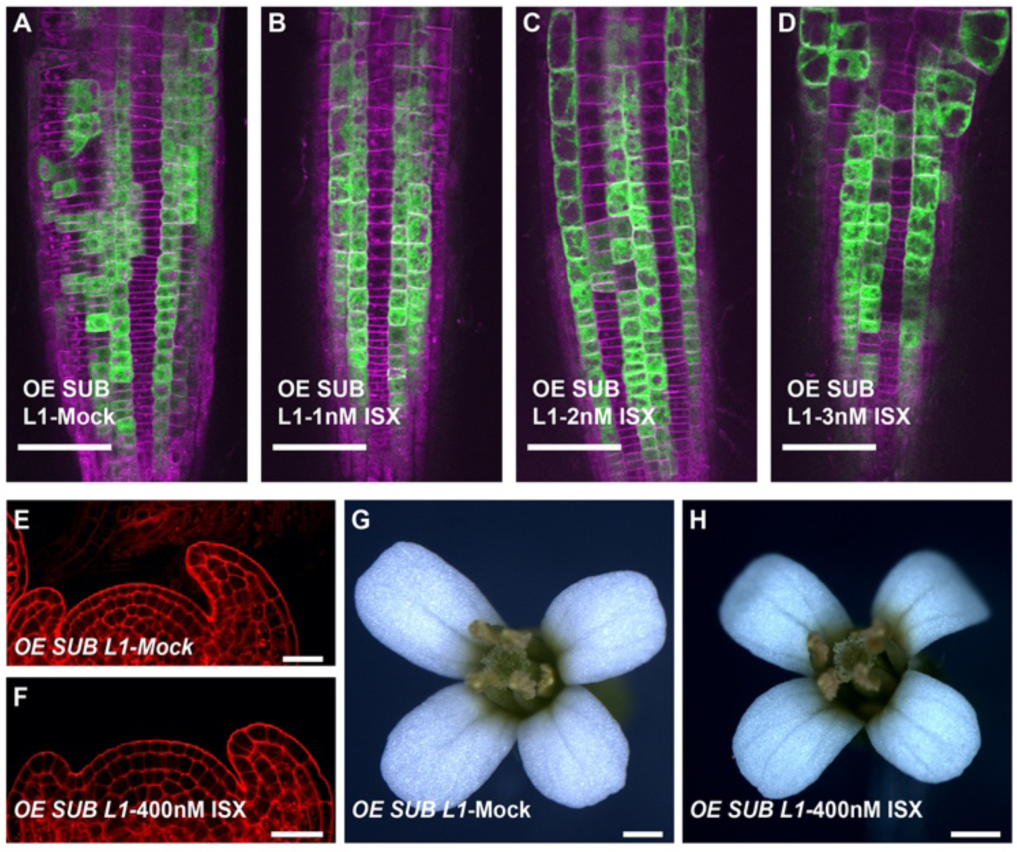
Overexpression of SUB:EGFP attenuates the isoxaben effects on root hair patterning and floral morphogenesis. (A-D) Confocal micrographs depict optical sections through roots of seven-day-old seedlings showing the pGL2::GUS:EGFP expression pattern. Genotypes and/or treatments are indicated. The cell wall was counter-stained with propidium iodide. (A) Note the mild aberration in pGL2::GUS:EGFP pattern (compare with Fig. 3H). (B-D) Note the normal to mildly abnormal pGL2::GUS:EGFP patterns (compare with Fig. 3I). For a quantification of the effects see Table 2. (A) 18/18 (B) 18/18 (C) 17/17 (D) 18/18 roots analyzed showed this pattern. The remaining roots showed the occasional single cell displacement in the reporter expression as also sometimes seen in wild type (Table 2). The experiment was performed three times with similar results. Scale bars: (A-D) 50μm, (E,F) 20μm, (G,H) 0.5mm.

Ectopic expression of *SUB* in *p35S::SUB* plants results in aberrant pGL2::GUS expression and a mild defect in root hair patterning (57). Confirming this finding, we found that L1 and O3 showed an altered expression pattern of the pGL2::GUS:EGFP reporter in roots of untreated seedlings, with more cells in the H position and fewer cells in the N position exhibiting reporter signal compared to wild type (Fig. 5A) (Table 2). In the case of the isoxaben-treated L1 and O3 lines we detected significant differences to wild type. Exposing seedlings of L1 and O3 to 1 or 2 nM isoxaben resulted in pGL2::GUS:EGFP patterns that resembled the pattern observed in untreated wild-type seedlings and that were less aberrant than the pGL2::GUS:EGFP patterns observed in corresponding isoxaben-treated wild-type seedlings (Fig. 5B-D) (Table 2). Moreover, the defects were weaker when compared to the aberrations exhibited by untreated L1 and O3. Exposing L1 or O3 seedlings to 3 nM isoxaben resulted in defects that were still less severe in comparison to wild-type plants treated with 3 nM isoxaben (Table 2).

Finally, we tested if ectopic expression of *SUB* also alleviates the effects of isoxaben on floral development by cultivating lines L1 and O3 in the presence of different concentrations of isoxaben as described above. We found that floral meristem and ovule defects were noticeably reduced in both lines compared to wild type (Fig. 5E-H) (Tables 3, 4).

## Discussion

The extracellular matrix in animal cells not only provides structural support but also exerts additional functions, including developmental patterning, as in the control of epidermal stem cell fate (17, 18). By contrast, the role of the plant cell wall in the regulation of cell fate is less well explored. The presented results strongly indicate that alterations in cell wall composition induced by the herbicide isoxaben or genetically in *prc1-1* mutants are associated with defects in root hair patterning in the root epidermis. Thus, they demonstrate a role of the plant cell wall in the control of cell fate.

The combined data indicate a shared molecular framework underlying cell-wall-mediated regulation of root hair cell fate, floral morphogenesis, and ovule development that involves the control of the atypical receptor kinase SUB, a well-known regulator of these developmental processes (19–21, 57). We found that exposure of seedlings to sub-lethal concentrations of isoxaben affects the architecture of SUB:EGFP-containing protein complexes at the plasma membrane, leads to increased internalization of SUB:EGFP, and causes a dose-dependent reduction in *SUB* transcript levels. The decrease of *SUB* activity is of biological consequence as ectopic expression of SUB:mCherry attenuated the detrimental effects of isoxaben on all the investigated developmental processes. Thus, the control of *SUB* activity by the cell wall is a central aspect of *SUB*-mediated signal transduction.

Our data expose a quantitative effect of cellulose reduction on *SUB* activity and concomitantly reveal that different levels of *SUB* activity are limiting for different *SUB*-dependent processes. We propose that the regulation of root hair patterning or floral morphogenesis depends on higher levels of *SUB* activity while the CBI-induced compensatory CWD response in seedlings requires only a basal level of *SUB* activity. This notion is supported by the observation that exposure of seedlings to sub-lethal doses of isoxaben resulted in reduced *SUB* transcript levels in seedlings and prominent defects in root hair patterning. Under such conditions the *SUB*-dependent CBI-induced CWD response remains operational (26).

Moreover, *QKY* is essential for *SUB*-mediated root hair patterning, floral morphogenesis, and ovule development (20, 23–25). However, with the exception of lignin accumulation, *QKY* is not required for the *SUB*-mediated CBI-induced CWD response (26) despite the fact that *SUB* levels are noticeably reduced in *qky* seedlings (25) (Fig. S2). Thus, changes in SUB complex architecture at the plasma membrane might underlie the different cell wall-dependent SUB functions. Interestingly, we observed a reduction in pSUB::SUB:EGFP signal in *prc1-1* while *SUB* transcript levels were unaltered. The finding suggests that a decrease in functional SUB protein levels is sufficient to lead to mild defects in root hair patterning. Further evidence for a limiting role for *SUB* activity in this process is provided by the observation that *sub-9* heterozygotes exhibit a *prc1-1*-like root hair patterning phenotype (Fig. 3M).

The differential effects of *PRC1* and isoxaben on *SUB* activity are consistent with a model that the control of *SUB* activity includes at least two distinct processes that react to variations in the CWD signal: post-transcriptional regulation and control of *SUB* transcript levels. A comparably low reduction in cellulose content would originate a CWD signal that leads to increased SUB internalization. A more pronounced drop in cellulose content would elicit a stronger or different CWD signal that would further affect *SUB* transcript levels. The stronger effect of higher concentrations of isoxaben in comparison to *prc1-1*, which carries a mutation in the *CESA6* gene (33), is explained by the observation that isoxaben affects several primary cell wall CESA subunits (29–31). Moreover, *CESA6* function is buffered by redundantly acting *CESA6*-like genes (6, 7).

There is crosstalk between cell wall components (12). For example, plants with a defect in hemicellulose production or in the pectin methylation exhibit reduced cellulose content (58, 59). Indeed, we found that *SUB* activity is downregulated upon application of epigallocatechin gallate (EGCG), an inhibitor of pectin methylesterase activity (60), to seven-day-old seedlings leading. Moreover, the treated seedlings exhibited root hair patterning defects (Fig. S3). The relative influence of altered pectin architecture and reduced cellulose content on these processes remains to be investigated.

The signal that controls *SUB* activity is currently unknown but the cell wall represents an obvious possible source. However, movements of CSCs are guided by cortical microtubules (8) and several components of CSCs interact with microtubules (61–66). Application of isoxaben results in the rapid internalization of CESA subunits (8) and isoxaben-treated wild-type plants as well as several *cesa* mutants were shown to exhibit altered cortical microtubule alignment (67–69). Thus, the signal regulating *SUB* activity could also originate from the cytoskeleton. However, we think it unlikely as treatment of seedlings with pharmaceutical compounds affecting the microtubule or actin cytoskeleton did not result in noticeable aberrations in root hair patterning. It will be interesting to identify the cell wall-derived signal in future studies.

## Materials and Methods

### Plant work and lines

Arabidopsis (L.) Heynh. var. Columbia (Col-0) and var. Landsberg (*erecta* mutant) (L*er*) were used as wild-type strains. Plants were grown on soil as described earlier (20). Plate-grown seedlings were grown in long-day conditions on half-strength Murashige and Skook (1/2 MS) agar plates supplemented with 0.3% sucrose. The following mutant alleles were used: *sub-1* and *qky-8* (L*er*) and *sub-9* and *sub-21* (Col) (20, 21, 26), *prc1-1* (33), *ixr2-1* (29). The lines carrying pSUB::SUB:EGFP, pUBQ::SUB:mCherry (O3, L1, Col), and pGL2::GUS:EGFP (Col) were reported in (24, 26–28). The pTHE1::THE1:GFP line was a gift from Herman Höfte. The pFER::FER:GFP (70) as well as the TMO7:1xGFP and TMO7:3XGFP reporter lines (39) were described previously. The generation of the various multiple mutant lines is detailed in SI, Materials and Methods.

### Chemical treatments

Isoxaben (ISX), 2,6-dichlorobenzonitrile (DCB), Thaxtomin A (TA), Latrunculin B (Lat-B), Cytochalasin D (Cyto-D), oryzalin, sorbitol, Congo Red, and epigallocatechin gallate (EGCG) were obtained from Sigma-Aldrich and used from stock solutions in DMSO (ISX: 100 µM, DCB: 1mM, TA: 100 µM, Lat-B: 100 µM, Cyto-D: 500 µM, Oryzalin: 100 µM, EGCG: 1mM) or in water (Sorbitol: 1M, NaCl: 2.5M, Congo Red: 1%). FM4-64 was purchased from Molecular Probes (2 mM stock solution in water). For FM4-64 staining seedlings were incubated in 4 μM FM4-64 in liquid 1/2 MS medium for 5 min prior to imaging.

### PCR-based gene expression analysis

For quantitative real-time PCR (qPCR) of *SUB* 35 to 40 seedlings per flask were grown in liquid culture under continuous light at 18 °C for seven days followed by treatment with mock or 600 nM isoxaben for eight hours or on plates (21 °C, long-day conditions). With minor changes, RNA extraction and quality control were performed as described previously (71). cDNA synthesis, qPCR, and analysis were done essentially as described (72). Primers are listed in Table S1.

### Microscopy

Confocal laser scanning microscopy, including colocalization analysis, scanning electron microscopy, and fixing and staining of floral meristems, were essentially performed as described earlier (20, 28, 73). A detailed description is provided in Supplementary Materials and Methods.

### Statistics

Statistical analysis was performed with PRISM8 software (GraphPad Software, San Diego, USA).

## Acknowledgments

We acknowledge Ramon Torres Ruiz and other members of the Schneitz lab for helpful discussion and suggestions. We thank Ramon Torres Ruiz for help with the fluorescence anisotropy analysis, Herman Höfte for the *prc1-1* allele and the pTHE1::THE1:GFP line, Martin Stegmann for the pFER::FER:GFP line, and Dolf Weijers for the TMO7:1xGFP and TMO7:3xGFP lines. We further acknowledge support by the Center for Advanced Light Microscopy (CALM) of the TUM School of Life Sciences. This work was funded by the German Research Council (DFG) through an Emmy Noether grant (WO 1660/2-2) to SW and an SFB924 grant (TP A2) to KS.

## Author Contributions

A.C., S.W. and K.S. designed research; A.C., X.C., B.L., J.G. and S.W. performed research;

A.C., X.C., B.L., J.G., S.W. and K.S. analyzed data; A.C., S.W. and K.S. wrote the paper.

## Supplementary Information

### Supplementary Figures

**Fig. S1.**
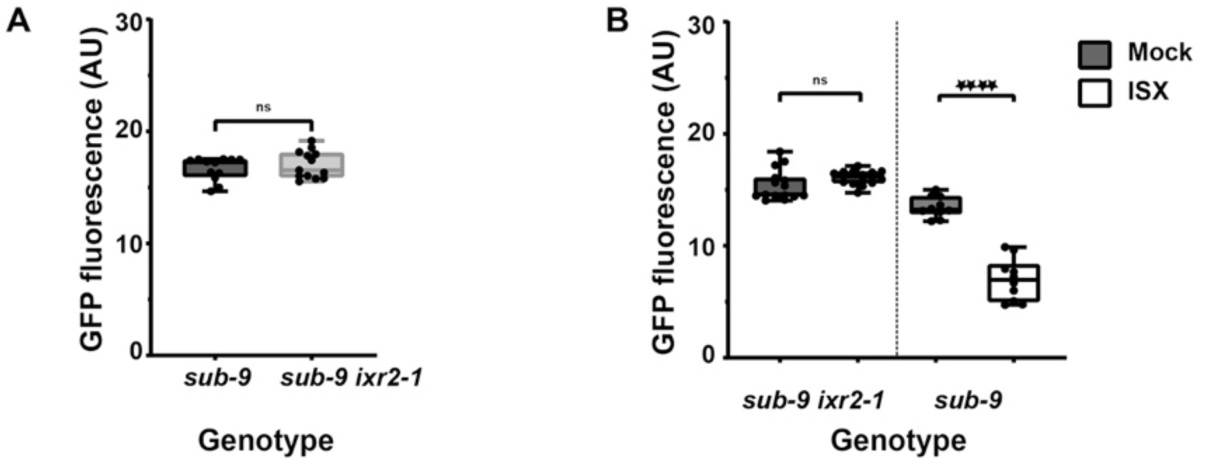
Effect of isoxaben treatment on pSUB::SUB:EGFP expression. (A) Quantification of signal intensity of the pSUB::SUB:EGFP reporter in seven-day-old *sub-9* and *sub-9 ixr2-1* plate-grown seedlings. Imaging parameters between both genotypes were identical. Box and whisker plots are shown. n ≤ 15. No statistical significance difference was observed (unpaired t-test with Welch’s correction, two-tailed P values). The experiment was performed three times with similar results. (B) Quantification of signal intensity of the pSUB::SUB:EGFP reporter in seven-day-old plate-grown *sub-9* and *sub-9 ixr2-1* seedlings transferred to plates containing 600 nM isoxaben for 8 hours. Imaging parameters between both genotypes were identical. Box and whisker plots are shown. 10 >n ≤ 15. No statistical significance difference was observed for treated or untreated sub-9 ixr2-1 seedlings (****, P < 0.0001; one-way ANOVA followed by post hoc Tukey’s multiple comparison test). The experiment was performed three times with similar results.

**Fig. S2.**
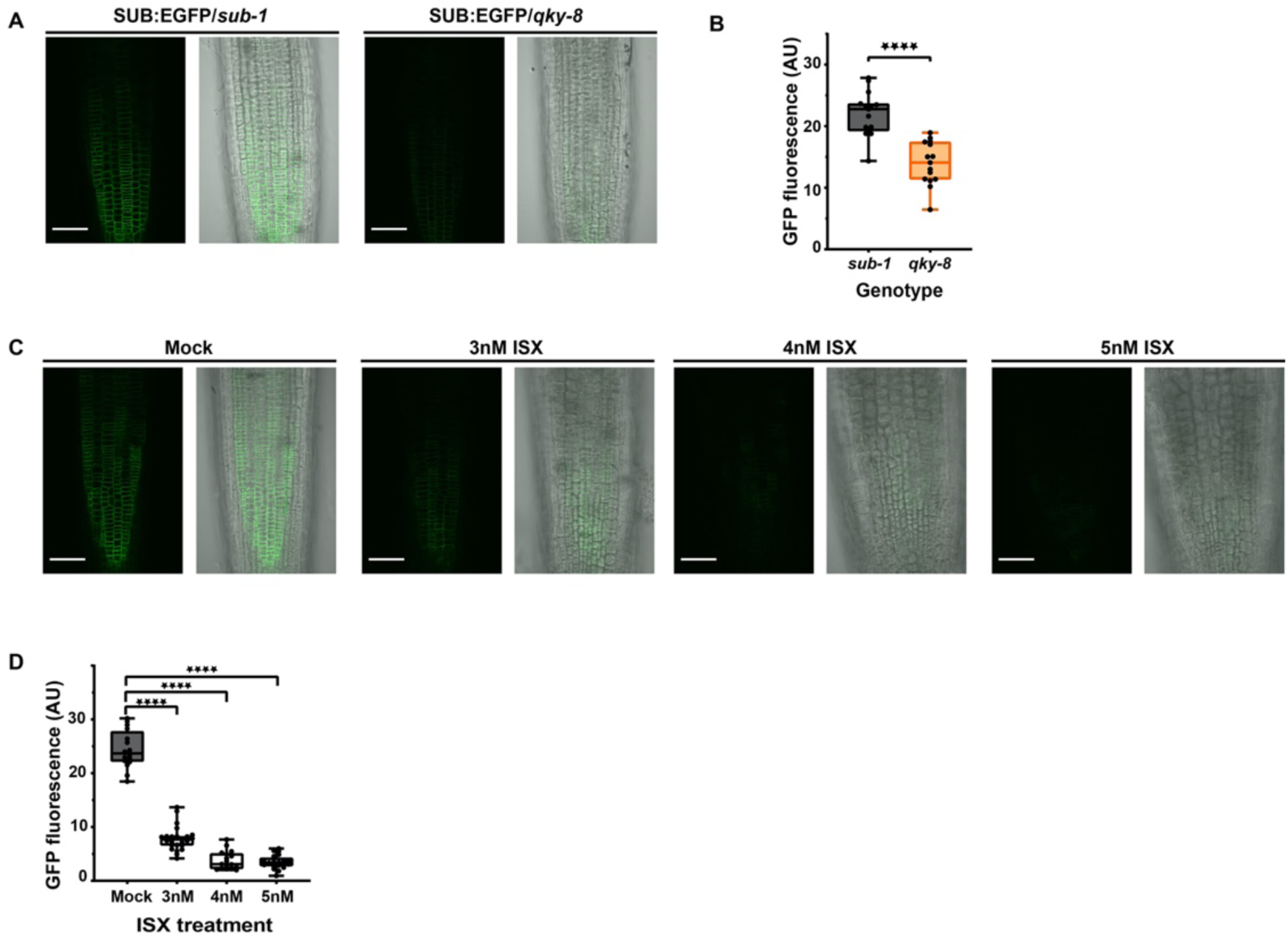
Effect of isoxaben treatment on pSUB::SUB:EGFP expression pattern in *qky-8*. (A,C) show confocal micrographs depicting optical sections through roots of seven-day-old seedlings. (A) Signal intensity of a functional pSUB::SUB:EGFP reporter in *sub-1* and *qky-8* genetic background. Note the reduction of signal in *qky-8*. (B) Quantification of the data shown in (A). Box and whisker plots are shown. n =15. Asterisks represent statistical significance (****, P < 0.0001; unpaired t-test with Welch’s correction, two-tailed P values). The experiment was performed three times with similar results. (C) Signal intensity of a functional pSUB::SUB:EGFP reporter in *qky-8*. Continuous treatments are indicated. Duration of treatment and treatment are indicated. Imaging parameters between the mock and isoxaben treatments were identical. Note the reduced signal in isoxaben-treated seedlings. (D) Quantification of the data shown in (C). Box and whisker plots are shown. 16 ≤ n ≤ 25. Asterisks represent statistical significance (****, P < 0.0001; one-way ANOVA followed by post hoc Tukey’s multiple comparison test). The experiment was performed three times with similar results. Scale bars: 50 μm.

**Fig. S3.**
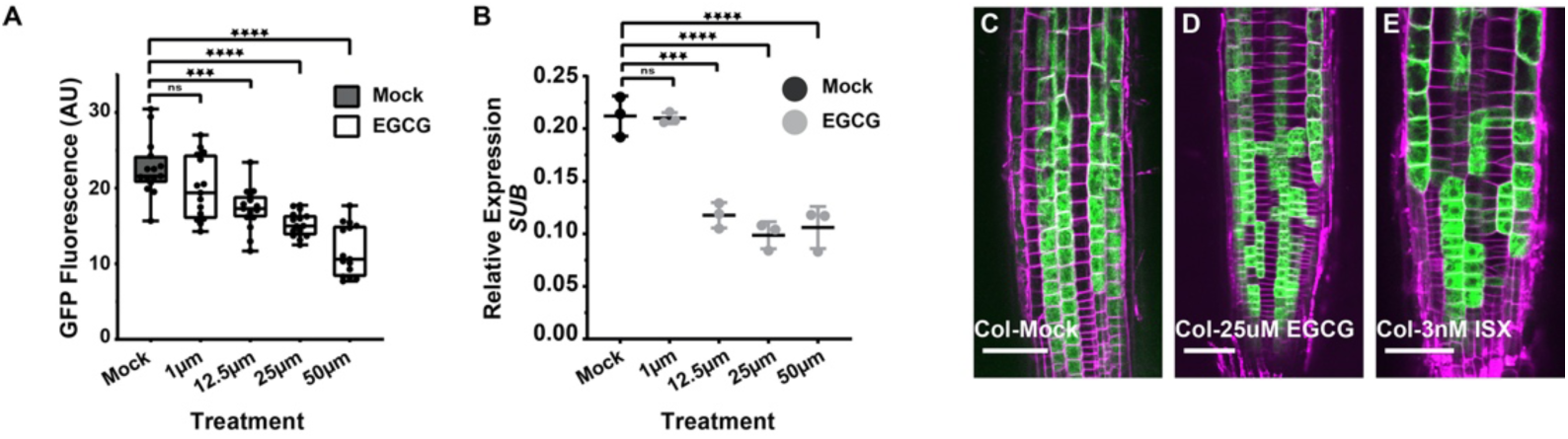
Effects of EGCG treatment on pSUB::SUB:EGFP, *SUB* and pGL2::GUS:EGFP expression patterns. (A) Box and whisker plot shows quantification of signal intensity of the pSUB::SUB:EGFP reporter upon continuous exposure of plate-grown seedlings to mock or to indicated amount of EGCG treatment (n=15). Asterisks represent statistical significance (***, P < 0.0006; ****, P < 0.0001; one-way ANOVA followed by post hoc Tukey’s multiple comparison test). The experiment was performed three times with similar results. (B) Relative transcript levels of *SUB* in seven-days-old seedlings exposed to mock or EGCG for indicated amount and time. Expression was detected by qPCR. Mean ± SD is shown, n=3. Asterisks represent statistical significance (***, P < 0.0006; ****, P < 0.0001; one-way ANOVA followed by post hoc Tukey’s multiple comparison test). The experiment was performed three times with similar results. (C-E) confocal micrographs depicting optical sections through roots of seven-day-old seedlings showing the pGL2::SUB:EGFP expression pattern. Genotypes and/or treatments are indicated. 10/10 (C), 13/15, (D) 11/14 (E) root analyzed showed this pattern. Scale bars: 50 μm.

### Supplementary Materials and Methods

#### Plant genetics

To generate the *sub-9 ixr2-1* double mutant F2 progeny of a parental cross between *sub-9* and *ixr2-1* were genotyped to select the double mutants. To generate the pSUB::SUB:EGFP sub-9 ixr2-1 and pSUB::SUB:EGFP sub-9 prc1-1 lines, the previously reported pSUB::SUB:EGFP plasmid was transformed into *sub-9*. Homozygous complementing T3 *pSUB::SUB:EGFP sub-9* was crossed into *sub-9 prc1-1* and *sub-9 ixr2-1* double mutants, respectively, and the F2 progeny was screened for double mutants with pSUB::SUB:EGFP expression and further propagated to obtain homozygous F3 lines. The *sub-9* pGL2::GUS:EGFP and *prc1-1* pGL2::GUS:EGFP lines were obtained by crossing the reporter line pGL2::GUS:EGFP into *sub-9* and *prc1-1*.

#### Microscopy

Confocal laser scanning microscopy was performed with an Olympus FV1000 set-up using an inverted IX81 stand and FluoView software (FV10-ASW version 01.04.00.09) (Olympus Europa GmbH, Hamburg, Germany) equipped with a water-corrected ×40 objective (NA 0.9) at ×3 digital zoom. For GFP fluorescence intensity measurements The mean grey values of GFP fluorescence signal intensity in the root epidermis of the pSUB::SUB:EGFP reporter was analyzed with ImageJ software (1). For each root, a region located 500 μm above the root tip (excluding the root cap) was used for analysis. To obtain the ratio between signal in cytoplasm versus total fluorescence intensity per cell by measuring signal intensity in two different regions of interests (ROIs). One ROI covered the cytoplasm and the other ROI included the cytoplasm and the outer cell boundary.

Steady state fluorescence anisotropy was measured using a FV3000 confocal laser scanning microscope (Olympus Europa GmbH, Hamburg, Germany) equipped with a single photon counting device with picosecond time resolution (LSM upgrade kit, PicoQuant, Berlin, Germany). GFP was excited at 485 nm with a linearly polarized, pulsed (40 MHz) diode laser (LDH-D-C-485, PicoQuant) using a 60x water immersion objective (Olympus UPlanSApo, NA1.2).The emitted light was collected in the same objective and was separated into perpendicular and parallel polarization with respect to excitation polarization. GFP fluorescence was then detected by an PMA Hybrid 40 detector (PicoQuant, Berlin, Germany) in a narrow range of its emission spectrum (BP520/35. A minimum 25 photons per pixel were counted with time correlated single photon counting (TCSPC) resolution of 25ps of ROI size 512px x 512px with 0.103 μM/px. Pixel-wise anisotropy was calculated with help of SymPhoTime 64 software (PicoQuant, Berlin, Germany) using instrument correction factors L1(0.035) and L2 (0.031) as well as the G-factor (1.018).

For SUB:EGFP subcellular localization upon drug treatments or colocalization with FM4-64, confocal laser scanning microscopy was performed on epidermal cells of root meristems located about 8–12 cells above the quiescent center using a Leica TCS SP8 X microscope equipped with GaAsP (HyD) detectors. The following objectives were used: a water-corrected ×63 objective (NA 1.2), a ×40 objective (NA 1.1), and a ×20 immersion objective (NA 0.75). Scan speed was set at 400 Hz, line average at between 2 and 4, and the digital zoom at 4.5 (colocalization with FM4-64), 3 (drug treatments), or 1 (root hair patterning). EGFP fluorescence excitation was performed at 488 nm using a multi-line argon laser (3% intensity) and detected at 502 to 536 nm. FM4-64 fluorescence was excited using a 561 nm laser (1% intensity) and detected at 610–672 nm. For the direct comparisons of fluorescence intensities, laser, pinhole, and gain settings of the confocal microscope were kept identical when capturing the images from the seedlings of different treatments. For determination of colocalization, the distance from the center of each EGFP spot to the center of the nearest FM4-64 signal was measured by hand on single optical sections using ImageJ/Fiji software (1). If the distance between two puncta was below the resolution limit of the objective lens (0.24 μm) the signals were considered to colocalize (2). Arabidopsis seedlings were covered with a 22×22 mm glass coverslip of 0.17 mm thickness (no. 1.5H, Paul Marienfeld GmbH & Co. KG, Lauda-Königshofen, Germany). Scanning electron microscopy was performed essentially as reported previously (3). Images were adjusted for color and contrast using ImageJ/Fiji software.

### Supplementary Tables

**Table S1.**
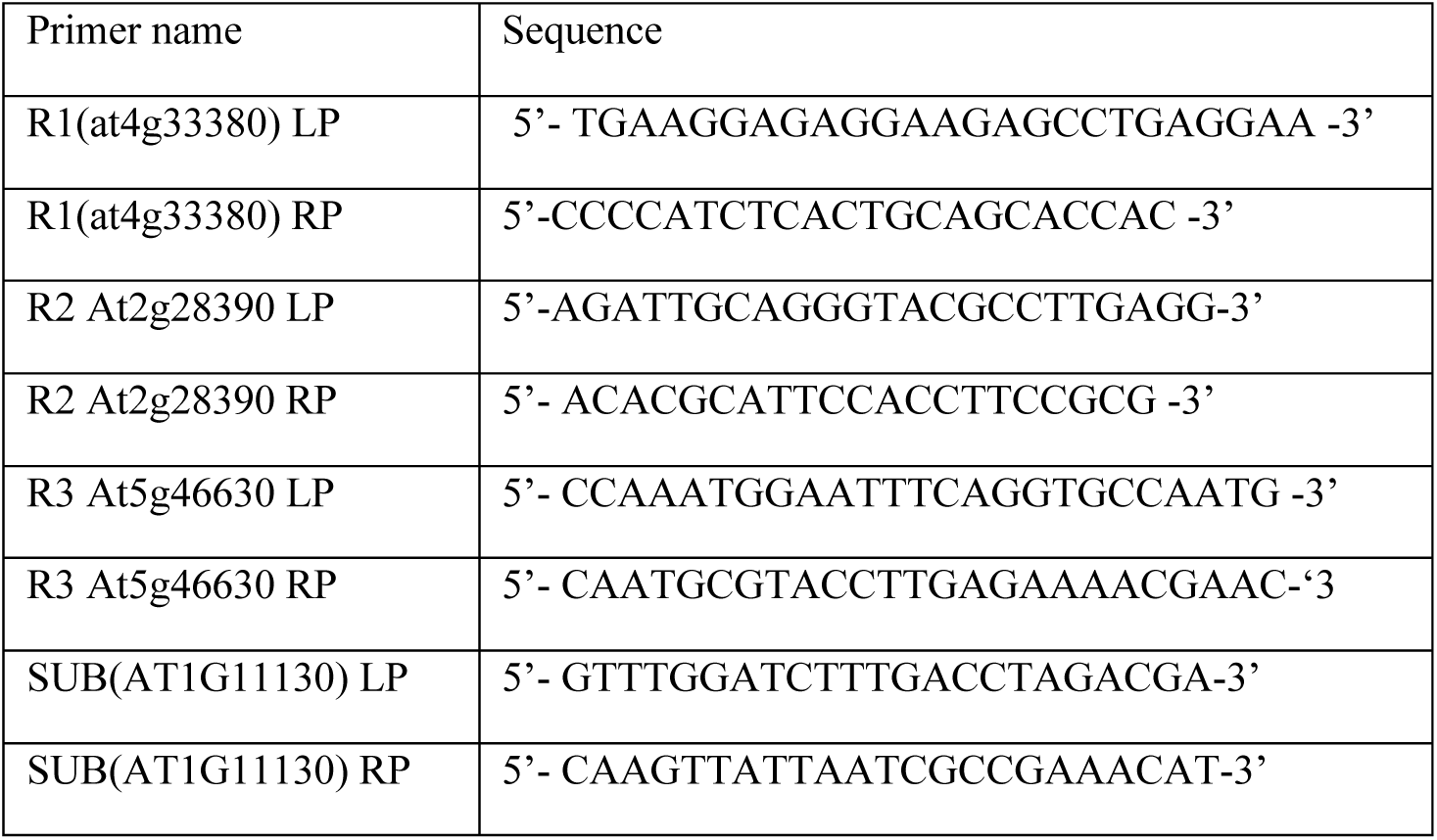
Primers used in this study

